# The brain dynamics of congenitally blind people seeing faces and letters via sound

**DOI:** 10.1101/2025.01.22.634358

**Authors:** Pawel J. Matusz, Lior Reich, Louise A. Stolz, Chrysa Retsa, David A. Tovar, Ella Striem-Amit, Elena Aggius-Vella, Amir Amedi, Micah M. Murray

## Abstract

Sensory substitution devices (SSDs) convert images to sounds to equip blind individuals with nominally visual functions, like face or letter sensitivity. Prior studies showed that image-to-sound SSDs engage cortices ordinarily specialised for visual functions. However, the brain dynamics of SSD-supported perception remains unknown. Either visual cortices are the first locus of discrimination of SSD percepts, or their activation is a byproduct of perceptual processes unfurling elsewhere, such as in auditory cortices. Resolving this uncertainty is critical for understanding the contribution of visual cortices to mechanisms subserving SSD-induced perceptions in the blind. Using electrical neuroimaging of EEG data from congenitally blind adults, we show for the first time that visual cortices are the earliest site of face and letter sensitivity when conveyed via SSDs, though with distinct temporal dynamics and spatial localizations. In the case of faces versus scrambled faces, differences first manifested at 460ms post-stimulus onset as topographic EEG modulations that were in turn localized to a network of right-hemisphere regions, including the fusiform face area (FFA) as well as lateral occipital cortices. In the case of discerning vertically versus horizontally oriented letter shapes, differences first manifested at 370ms post-stimulus onset as topographic EEG modulations that were localized to a network of left-hemisphere regions, including the occipital pole and the occipito-temporal junction extending into the angular gyrus. Notably, early-latency auditory evoked potentials did not differ based on visual properties of the soundscapes. These collective data support the proposition that responses to soundscapes are routed to brain circuits ordinarily specialized for processing visual information and that these circuits are the loci of the initial discriminant processes. By providing the temporal dynamics of SSD perception, our findings provide unique evidence for the theory that cortices are characterised by task-contingent functional organisation.

**Highlights:**

- Sensory substitution devices (SSDs) provide the blind access to visual information
- Brain dynamics in visual cortices to SSD-conveyed object images are unknown
- Early EEG sensitivity for faces and letters was localized to the visual cortex
- The blind visual cortex takes part in early SSD processing for visual categories

**eTOC blurb:** We characterise for the first time the brain dynamics of the blind using soundscapes for face and letter discrimination. Faces recruited the FFA at 460ms post-stimulus onset, whereas letters recruited the occipital pole, occipito-temporal junction, and angular gyrus at 370ms post-stimulus onset. Our findings of the temporal dynamics of SSD perception uniquely support the theory that cortices are characterised by task-selective sensory-independent functional organisation.

## Introduction

Visual-to-auditory sensory-substitution devices (SSDs) empower the blind to harness auditory information to perform nominally visual functions, like navigation or object recognition. Notably, the SSD-entrained sound stimuli can engage the same brain networks as those activated during visual processing in sighted individuals^1–3^. This highlights the brain’s natural capacity for plasticity in the service of goal-directed behavior. In the absence of visual sensory inputs, the brain adapts to alternative inputs. A critical question is whether the blind truly use their visual cortex to process this information. Brain imaging and mapping methods such as fMRI and EEG are well-poised to address this. While fMRI can delineate *where* SSD-induced vision transpires, it cannot resolve the underlying brain dynamics of where and *when* these processes first occur. Either visual cortices are the initial locus of discrimination of SSD percepts, or their activation is a byproduct of perceptual processes unfurling first elsewhere. Resolving this uncertainty is critical for understanding whether and under which circumstances the blind may truly process SSDs with their visual cortices.

For both blind and sighted participants alike, auditory inputs delivered via auditory-to-visual SSDs enable high-level visual functions. Psychophysical studies have shown that SSDs equip people with skills in recognizing object categories as well as specific object features, e.g., spatial location, size, shape, texture, and orientation^4–7^. This evidence in turn motivated the use of neuroimaging to identify the nature of processes underlying the SSD-enabled skills. If the entrained sounds first engage the same visual cortices as those typically observed in sighted individuals, this would be evidence that visual cortices can be repurposed for non-visual processing in the blind when using SSDs. In turn, if SSD-induced perceptual discrimination first occurs outside of visual cortices, this would be more in line with the blind’s visual cortices’ responses as dissimilar from its typical function in the sighted brain; possibly to reinforce functions such as acoustic discrimination or associative learning.

Prior fMRI research has provided evidence for SSD-induced activation of visual cortices^6,8–11^. For example, regions of the lateral occipital complex (LOC) that are typically involved in visual shape and object processing are also activated in response to SSD sounds of the same stimuli^9^. Regions of the right fusiform gyrus, including the fusiform face area (FFA), have similarly been shown to respond to face stimuli delivered as soundscapes^12^. Regions of the left fusiform gyrus, including the visual wordform area (VWFA), have been shown to respond to letter stimuli delivered as soundscapes^6^. Still other data show that the large-scale functional organization of the visual cortices, as exemplified by dissociation between object and spatial processing, is also observed when the congenitally blind are trained to use SSD sounds^13^. More importantly, this work has shown that such activations are dependent on training; there were no responses in these areas when the participants were unfamiliar with the SSDs^9,14^.

This series of works provide the first line of support that SSD-induced activity in these visual areas is linked to perception. Collectively, this evidence has been formalized in the hypothesis that the brain is a *task-contingent machine*, whereby brain areas are characterized by their predominant function rather than the sensory modality inputs they receive^15,16^. This task-contingent nature of the brain seems so strong, that several hours to weeks of training suffice for brain regions to adapt to a new type of sensory inputs as a source of task-relevant information. To support this task-contingent machine hypothesis more fully, it is necessary to ascertain if visual areas are themselves performing the key perceptual discrimination prior to other brain areas. Presently, this cannot be fully addressed with hemodynamic imaging, but rather requires methods that are time-resolved, such as M/EEG.

The present study applied electrical neuroimaging analyses to high-density EEG to characterize the brain dynamics of face and letter sensitivity conveyed via SSDs in a group of congenitally blind individuals who successfully trained with the vOICe algorithm^17^ (see also Methods for results of training). Electrical neuroimaging refers to a pipeline including analyses of response strength and topography as well as time-resolved source reconstructions^18–20^. The vOICe converts pixelated images to sounds by representing vertical position of a pixel by pitch (low to high) and horizontal position of a pixel by time (sweeping left to right). After training and during EEG recordings, participants passively listened to soundscapes of faces, scrambled faces, as well as at letter stimuli (stimulus duration = 1570ms). We analyzed event-related potentials (ERPs) elicited by soundscapes representing line drawings of faces and their scrambled counterparts, as well as Hebrew letters (**Figure 1**). Line drawing stimuli are particularly well-suited for conversion by the vOICe algorithm. Because the soundscapes represented a specific, well-studied object category (i.e. faces and letters), we had the advantage of clear predictions regarding the visual regions putatively involved; notably the inferotemporal cortices, including the FFA^21,22^ in the case of faces and VWFA^23^ as well as angular gyrus (AG)^24^ in the case of letters. Specifically, we hypothesized that responses to soundscapes of faces, but not those to scrambled faces, would engage the inferotemporal cortices (see ^12^ for comparable fMRI data in blind participants). We also hypothesized that the VWFA and AG would exhibit sensitivity to subgroups of soundscapes of letters (see ^6^ for comparable fMRI data in blind participants). Moreover, the timing of these activations, relative to any differential responses in auditory cortices, was crucial for our research questions. To our knowledge, the only prior EEG study of SSD training showed that the earliest differential responses to SSD stimuli were first in auditory cortices^25^, which runs counter to the task-contingent machine hypothesis. Additionally, they used geometric shapes as stimuli, and their sample consisted of sighted participants. As such, while their data indeed demonstrate brain plasticity following SSD training, the pertinence of their findings to the question of the nature of visual perceptions afforded by SSD in the blind is limited.

**Figure 1.**
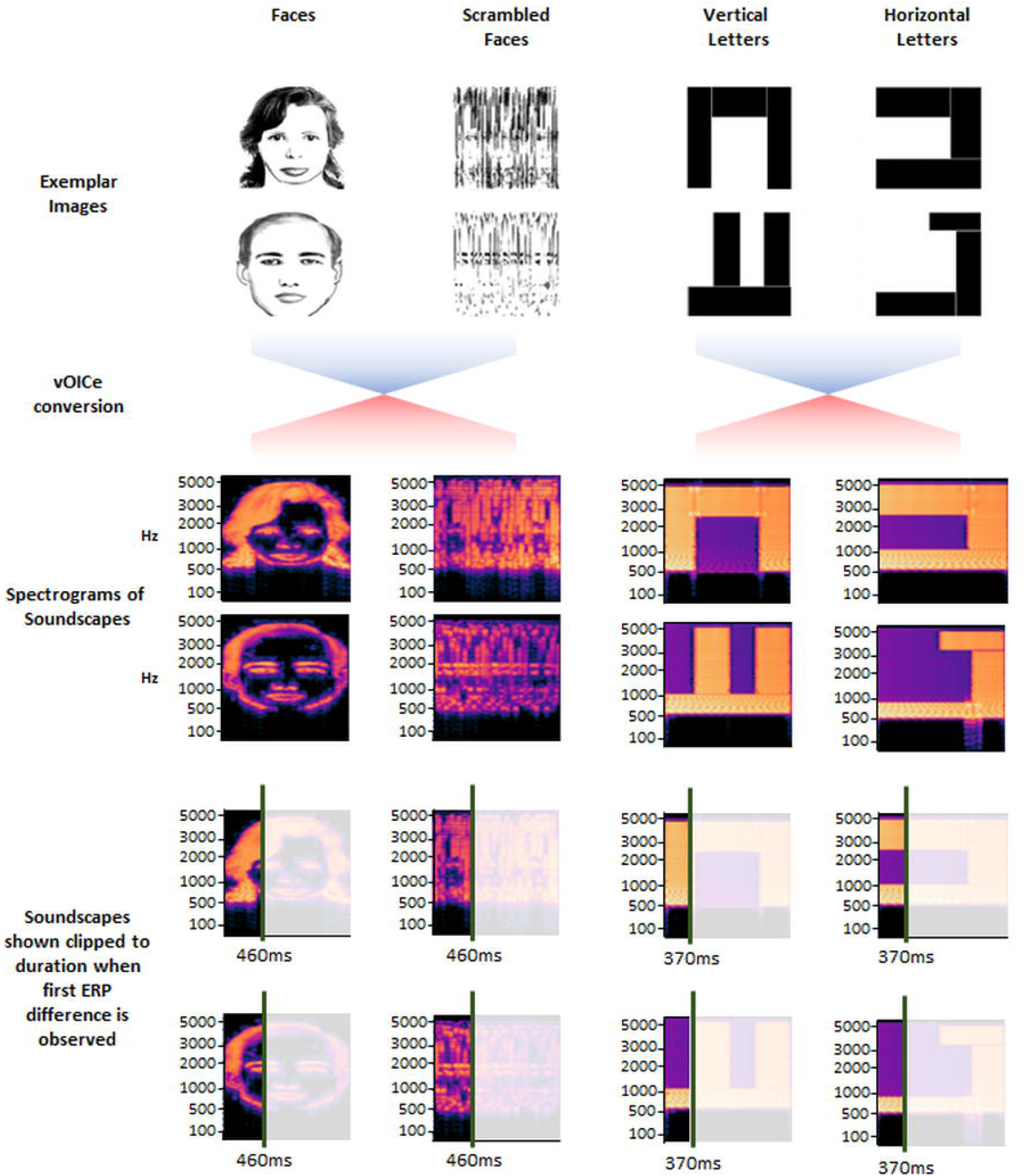
Image stimuli and spectrograms of soundscapes. Images included line drawings of faces and scrambled faces as well as vertically and horizontally oriented letters, examples of which are displayed. The middle panels illustrate how, after conversation via the vOICe algorithm, soundscapes retain image properties in their spectrogram. The lower panels show the same spectrograms clipped to the latency when ERP differences were first observed in the present study.

## Results

### Brain dynamics of SSD-enabled face sensitivity

For demonstrative purposes, we present the ERPs to the faces versus scrambled faces for three midline electrodes (**Figure 2A**). To determine the brain dynamics by which soundscapes of faces and scrambled faces are first discriminated, we first conducted a mass univariate analysis across both the peri-stimulus period (i.e., 100ms pre-stimulus to 900ms post-stimulus onset) and the entire 64-channel scalp electrode montage. This analysis entailed a non-parametric paired randomization test with 1000 shuffles at each sample as well as FDR correction for multiple comparisons. To further account for spatio-temporal autocorrelation, we only considered as reliable those effects that were temporally sustained (i.e., ≥10 consecutive data points^26^; see also^27–29^. By such criteria, no response differences were observed at the level of individual electrodes. Of note, there was no evidence of differential responses to face and scrambled face soundscapes during post-stimulus time periods of the auditory evoked potential (i.e. the P50-N1-P2 complex) that have been shown in other works to be modulated by differences in low-level acoustic parameters and to involve prominent sources along the superior temporal gyrus^30,31^.

**Figure 2.**
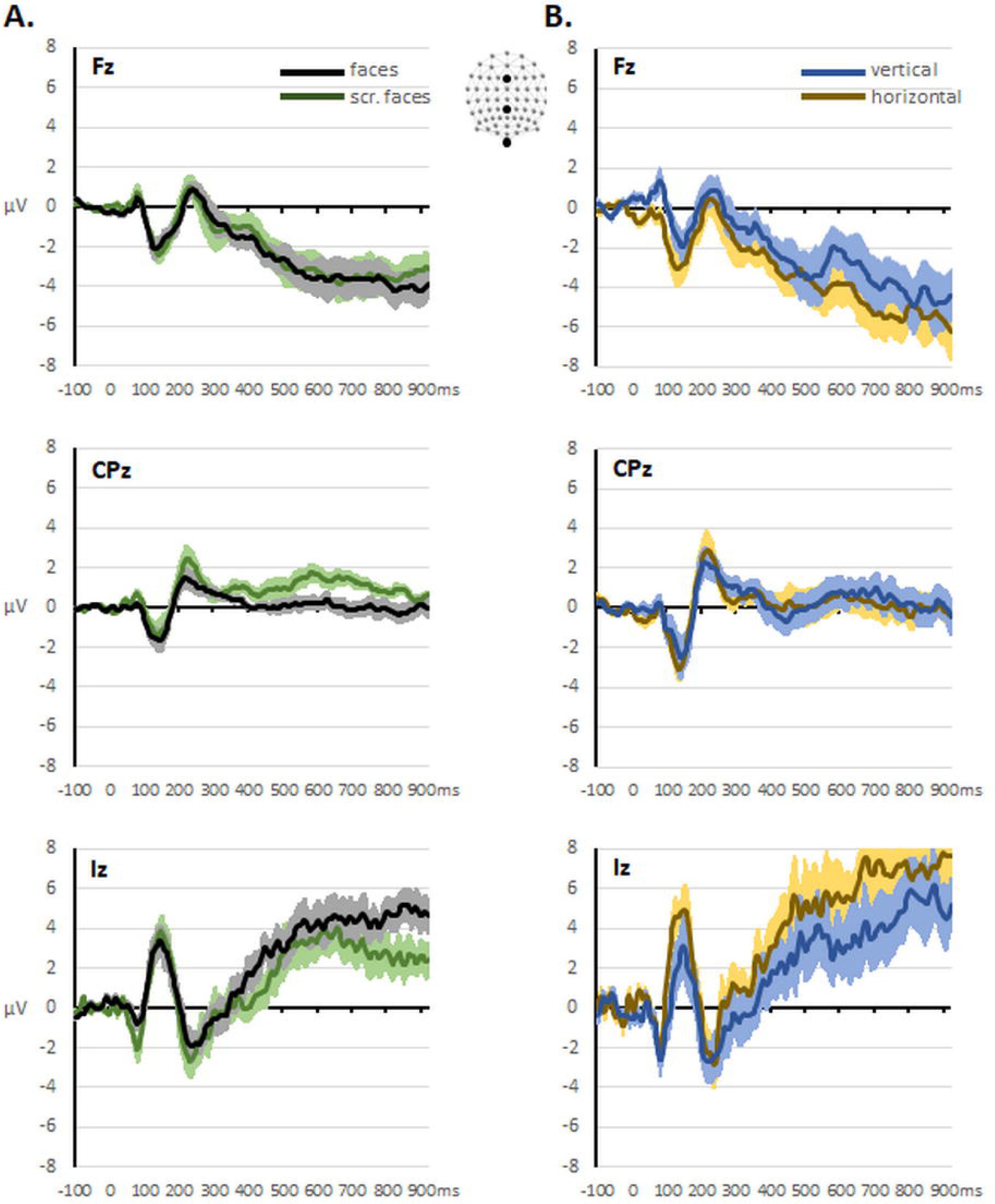
Exemplar group-averaged voltage ERP waveforms at a sample of midline scalp locations (see inset). (A) Mean (±SEM) voltage ERP waveforms in response to soundscapes of faces (black traces) and soundscapes of scrambled faces (green traces). (B) Mean (±SEM) voltage ERP waveforms in response to soundscapes of vertically oriented letters (blue traces) and soundscapes of horizontally oriented letters (brown traces). The mass univariate analyses failed to reveal sustained significant differences after FDR correction for multiple comparisons.

Next, we performed multivariate analyses on reference-independent measures of the ERP across the full 64-channel montage; namely the ERP strength, as quantified by Global Field Power, and modulations in the ERP topography as quantified by Global Dissimilarity^18,20^. For both metrics, analyses were conducted across the peri-stimulus period, as detailed above, and included a contiguous temporal criterion of ≥10 samples for an effect to be considered reliable. There was no evidence for statistically reliable modulations in response strength between conditions. By contrast, there was evidence for statistically significant topographic modulations over the 484-756ms period as well as 804-852ms period (**Figure 3A**). This multivariate analysis indicates that the earliest detectable ERP modulation followed from a change in the topography of the electric field at the scalp, which biophysical laws dictate must follow from changes in the underlying configuration of active sources in the brain^18^. In other words, different brain networks subserved responses to soundscapes of faces versus scrambled faces.

**Figure 3.**
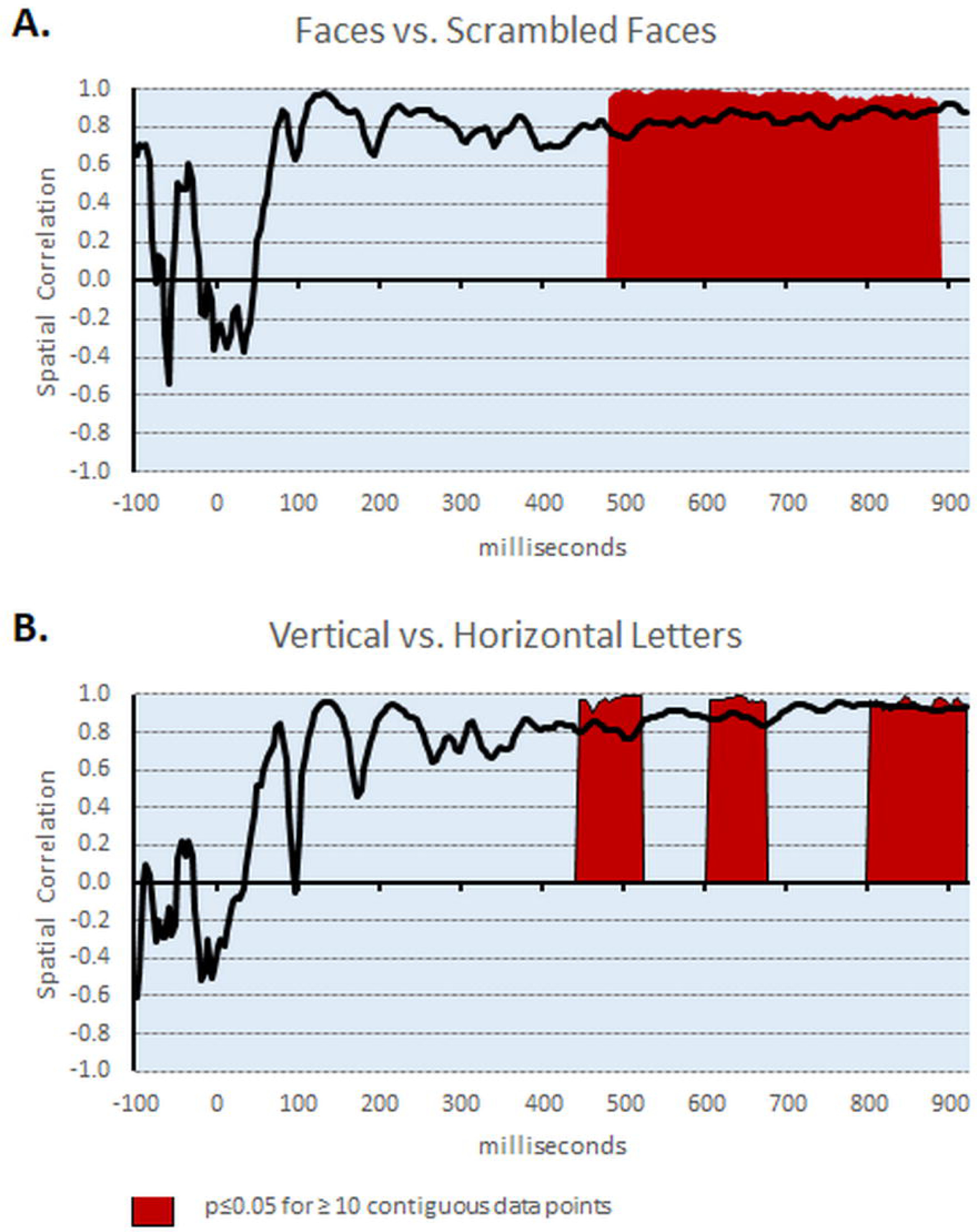
Results of multivariate analyses of topography of the electric field at the scalp. (A) Results from the analysis of soundscapes of faces and scrambled faces indicate topographic differences starting at 480ms post-stimulus onset (red area shows 1 minus p-value for timepoints exhibiting a significant difference). The black trace displays the spatial correlation between responses to soundscapes of faces and scrambled faces. (B) Corresponding results of the comparison of responses to vertically and horizontally oriented letters are displayed using the same conventions as is (A). Topographic differences onset at 444ms post-stimulus.

To better characterize these brain networks and their temporal stability, we next performed a hierarchical clustering analysis of the ERP topography, using as input the concatenated group-averaged data in response to soundscapes of faces and scrambled faces. The topographic clustering indicated that 7 different template maps accounted for 97.91% of the global explained variance of the dataset (**Figure 4A**). Until 456ms post-stimulus onset, the same sequence of four topographic patterns were observed in the group-average responses to both soundscapes of faces and scrambled faces. This sequence included topographies consistent with the P50-N1-P2 complex of auditory ERP components^30^. Therefore, the visual categorical information was not processed in the auditory cortex in these early time windows. However, over the 460-716ms period, the topographies of ERP responses in those conditions started to differ (**Figure 4A**). Specifically, one topography, framed in red, appeared to better characterize responses to soundscapes of faces, whereas another topography, framed in gray, appeared to better characterize ERP responses to soundscapes of scrambled faces. This was statistically tested by performing a competitive spatial correlation between each template map observed at the group-average level and the single-subject data from both conditions. This procedure yields a measure of how many time samples over the 460-716ms period exhibited a higher spatial correlation with each template map, which was in turn submitted to a Wilcoxon Signed Ranks test (p=0.015). On average, the template map framed in red better characterized responses to soundscapes of faces than scrambled faces (74.9±11.9% vs. 31.1±13.1%) and another map, framed in dark gray, better characterized responses to soundscapes of scrambled faces than faces (25.1±11.9% vs. 68.9±13.1%) (**Figure 4B**). This observation was also statistically tested for each individual participant using a Chi-square test comparing the distribution of the number of time samples characterized by each template map for responses to soundscapes of faces (actual distribution) with the distribution observed for responses to soundscapes of scrambled faces (expected distributions). Statistically significant differences were observed for 7 of the 9 participants (all p’s ≤0.01; **Figure 4B**). Distinct topographies characterizing responses to soundscapes of faces versus scrambled faces was also observed over the later 720-900ms post-sound onset period (**Supplementary Figure S2**), though here we focus on the earliest differences.

**Figure 4.**
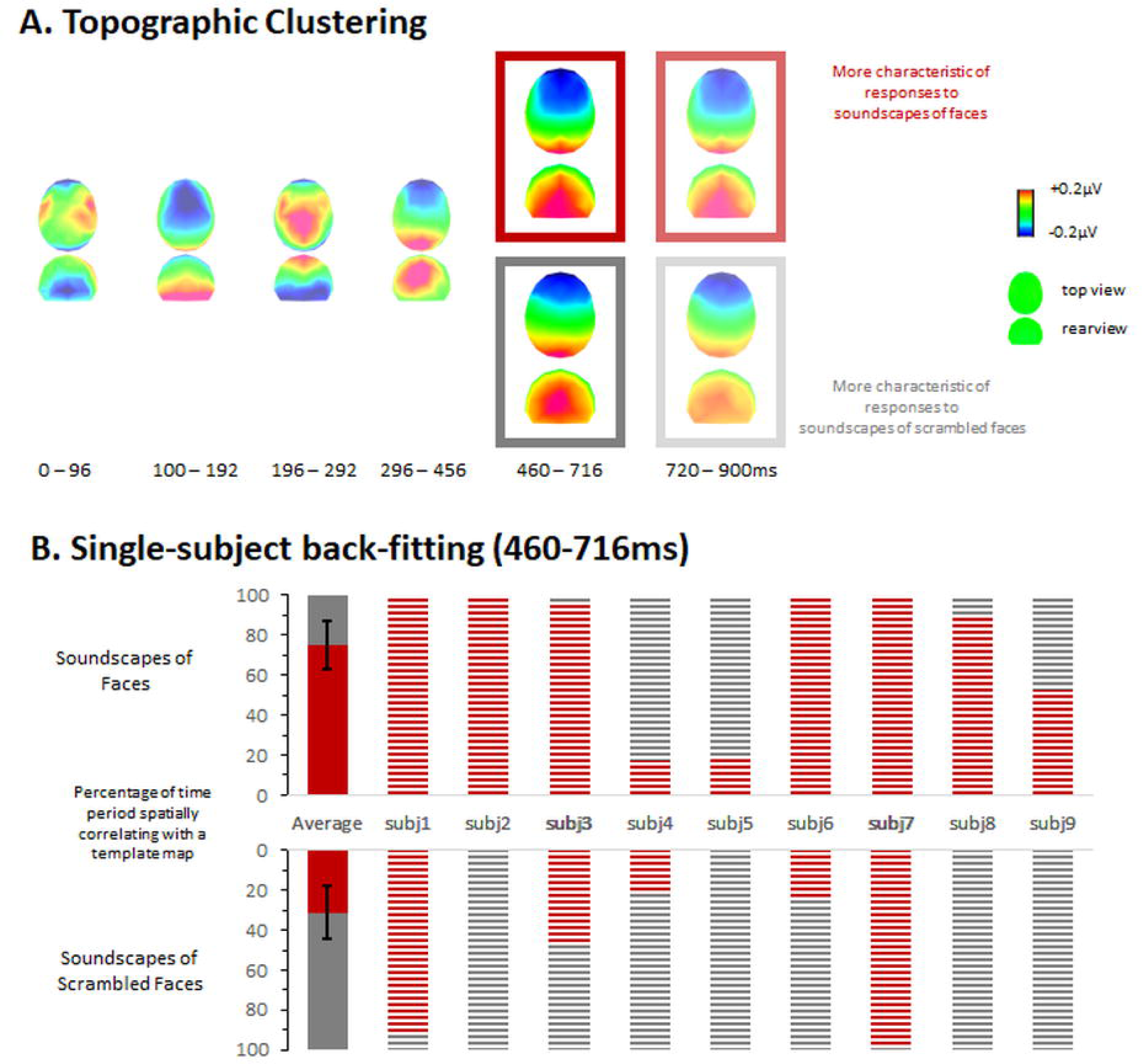
Topographic clustering results for responses to soundscapes of faces and scrambled faces. (A) The group-averaged ERPs were submitted to hierarchical clustering. Over several peristimulus onset time windows from 0 to 456ms identical ERP topographies were observed across both conditions. Over the 460-716ms post-stimulus time period, distinct maps appeared to characterize responses to face and scrambled face soundscapes (red- and gray-framed topographies, respectively). (B) This observation was confirmed both at the group-average and single-subject levels by backfitting template maps observed at the group-averaged level to single-subject data based on spatial correlation. The bar graph shows the group-averaged results (solid bars) as well as data from each participant (patterned bars). Similar differences were observed over the 720-900ms post-stimulus time period (see Supplementary Figure S2).

Data from the 460-716ms post-sound onset time were then submitted to EEG source modelling (**Figure 5**). The statistical contrast revealed significantly stronger (p<0.05 for >10 contiguous solution points) responses to soundscapes of faces than scrambled faces predominantly within the right inferotemporal cortices as well as within the junction between the right superior temporal cortex and insula (**Figure 5**). Within the inferotemporal cortices, there were 2 clusters: one with a local maximum in the right fusiform gyrus (Talairach^32^ coordinates: 27, −42, −11mm) and the other with a local maximum within the right lateral occipital cortex (42, −72, −4mm). A third cluster was observed within the right insula, extending into the right superior temporal cortex (maximum at 42, −4, −4mm). These results indicate that higher-order visual cortices are involved in the initial discrimination of soundscapes of faces versus scrambled faces.

**Figure 5.**
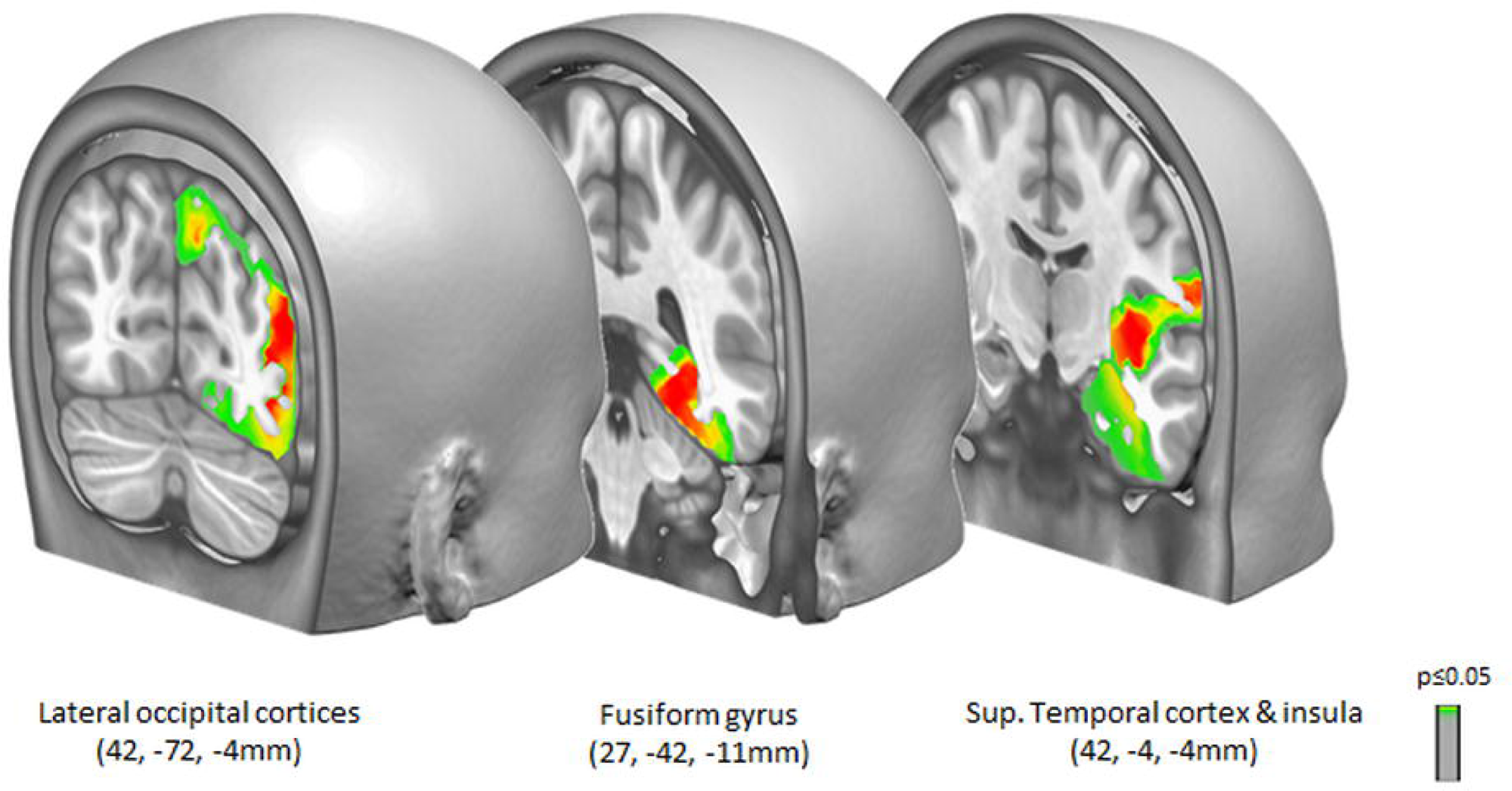
Source modelling analyses for responses to soundscapes of faces and scrambled faces. Data from the 460-716ms post-sound onset time were then submitted to distributed linear source modelling. Stronger responses (p<0.05 for >10 contiguous solution points) were observed in three clusters; the Talairach^32^ coordinates of the local maximum are indicated.

### Brain dynamics of SSD-enabled orientation sensitivity

Similarly to what is shown for responses to soundscapes of faces and scrambled faces, we likewise display voltage ERP waveforms in response to soundscapes of letters oriented vertically and horizontally from the same midline electrodes (Figure 2B). The mass univariate analysis across both the peri-stimulus period (i.e., 100ms pre-stimulus to 900ms post-stimulus onset) and the entire 64-channel scalp electrode montage failed to reveal significant differences (1000 shuffles and FDR correction). As such, there was again no evidence of differential responses at the single-electrode level during post-stimulus time periods of the auditory evoked potential (i.e. the P50-N1-P2 complex).

Multivariate analyses of the reference-independent metrics of response strength and response topography were subsequently performed. There was no evidence for differences in response strength as quantified from the GFP. By contrast, there were significant topographic differences observed over the 424-512ms and 608-660ms post-stimulus time periods (**Figure 3B**). This multivariate analysis indicates that the earliest detectable ERP modulation followed from topographic changes in the electric field at the scalp, which forcibly derive from changes in the underlying configuration of active sources in the brain. In other words, different brain networks subserved responses to soundscapes of letters with a predominant vertical versus horizontal orientation.

To better characterize these brain networks and their temporal stability, we next performed a hierarchical clustering analysis of the ERP topography, using as input the concatenated group-averaged data in response to soundscapes of vertical and horizontal letters. The topographic clustering indicated that 8 different template maps accounted for 98.1% of the global explained variance of the dataset (**Figure 6A**). Until 370ms post-stimulus onset, the same sequence of three topographic patterns were observed in the group-average responses to both soundscapes of faces and scrambled faces. This sequence included topographies consistent with the P50-N1-P2 complex of auditory ERP components, again showing that the auditory cortex dynamic response does not carry visual-like shape information for letters. Commencing at 370ms post-stimulus onset, the topographies of ERP responses in those conditions started to differ. Specifically, one topography appeared to better characterize responses to soundscapes of vertical letters, whereas another topography appeared to better characterize ERP responses to soundscapes of horizontal letters. This was statistically tested by performing a competitive spatial correlation between each template map observed at the group-average level and the single-subject data from both conditions. This procedure yields a measure of how many time samples over the 370-520ms period exhibited a higher spatial correlation with each template map, which was in turn submitted to a Wilcoxon signed rank test (p=0.046). On average, the template map framed in blue better characterized responses to soundscapes of vertical than horizontal letters (58±13.9% vs. 29.2±13.1%) and another map, framed in brown, better characterized responses to soundscapes of horizontal than vertical letters (70.8±13.1 vs. 41.9±13.9%) (**Figure 6B**). This observation was also statistically tested for each individual participant using a Chi-square test comparing the distribution of the number of time samples characterized by each template map for responses to soundscapes of vertical letters (actual distribution) with the distribution observed for responses to soundscapes of horizontal letters (expected distributions). Statistically significant differences were observed for 5 of the 9 participants (all p’s ≤0.05; **Figure 6B**). Distinct topographies characterizing responses to soundscapes of vertical versus horizontal letters was also observed over the later 524-900ms post-sound onset period (**Supplementary Figure S3**), though here we focus on the earliest differences.

**Figure 6.**
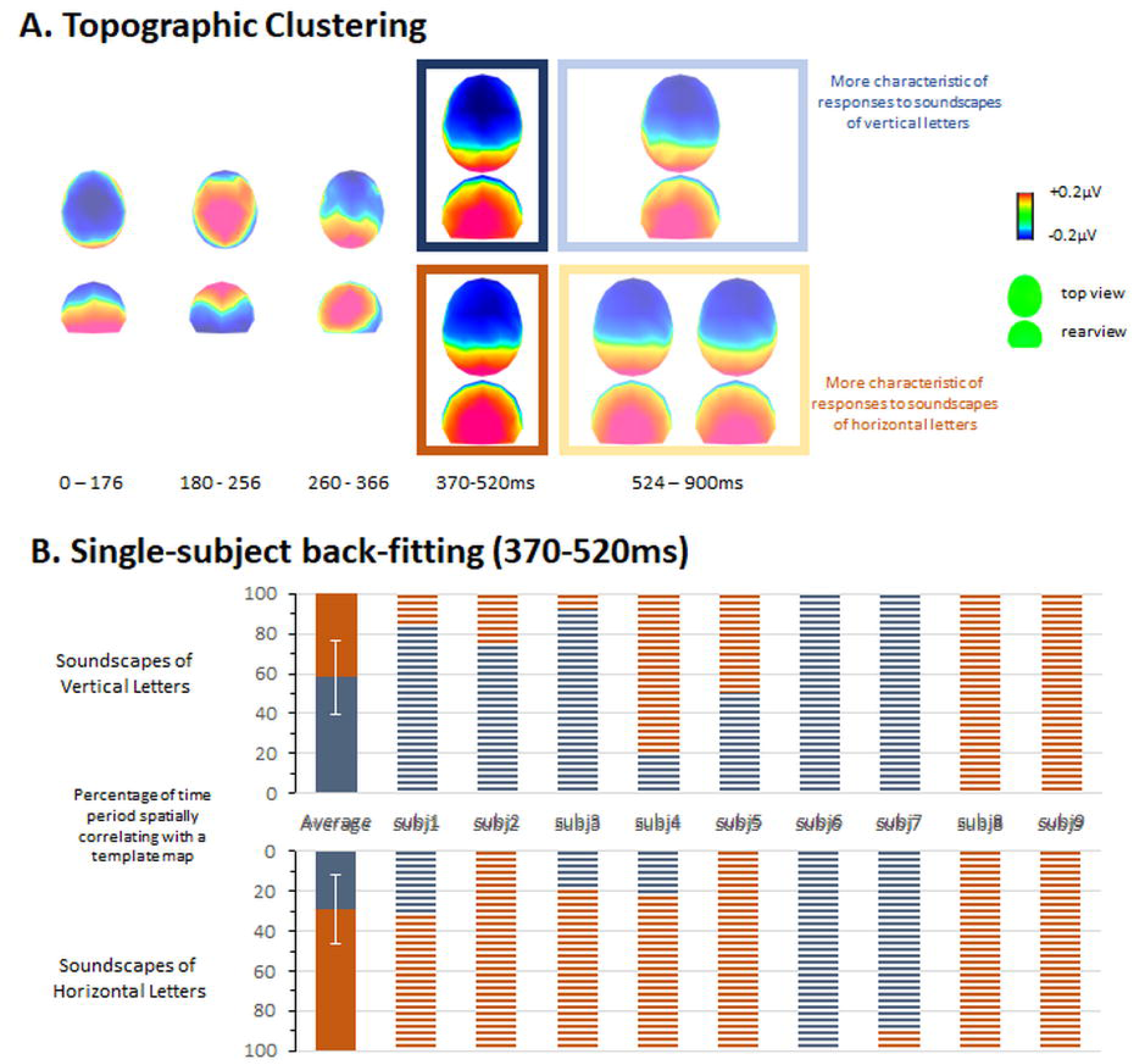
Topographic clustering results for responses to soundscapes of vertically and horizontally oriented letters. (A) The group-averaged ERPs were submitted to hierarchical clustering. Over several peristimulus onset time windows from 0 to 366ms identical ERP topographies were observed across both conditions. Over the 370-520ms post-stimulus time period, distinct maps appeared to characterize responses to vertical and horizontal letter soundscapes (blue- and brown-framed topographies, respectively). (B) This observation was confirmed both at the group-average and single-subject levels by backfitting template maps observed at the group-averaged level to single-subject data based on spatial correlation. The bar graph shows the group-averaged results (solid bars) as well as data from each participant (patterned bars). Similar differences were observed over the 524-900ms post-stimulus time period (see Supplementary Figure S3).

Data from the 370-520ms post-sound onset time were then submitted to EEG source modelling (**Figure 7**). The statistical contrast revealed significantly stronger (p<0.05 for >10 contiguous solution points) responses to soundscapes of vertical than horizontal letters within the left posterior-inferior occipital cortex (Talairach^32^ coordinates: −33, −85, −11mm) and left occipito-temporal cortices (−49, −62, 13mm) that extended anteriorly and superiorly into the angular gyrus (−49, −61, 34mm). Significantly stronger responses to soundscapes of horizontal than vertical letters were observed within the right inferior parietal cortex (64, - 24, 32mm). These results indicate that visual cortices which have been implicated in processing of visual letters and words are also the first involved in the initial discrimination of soundscapes of vertical versus horizontal letters.

**Figure 7.**
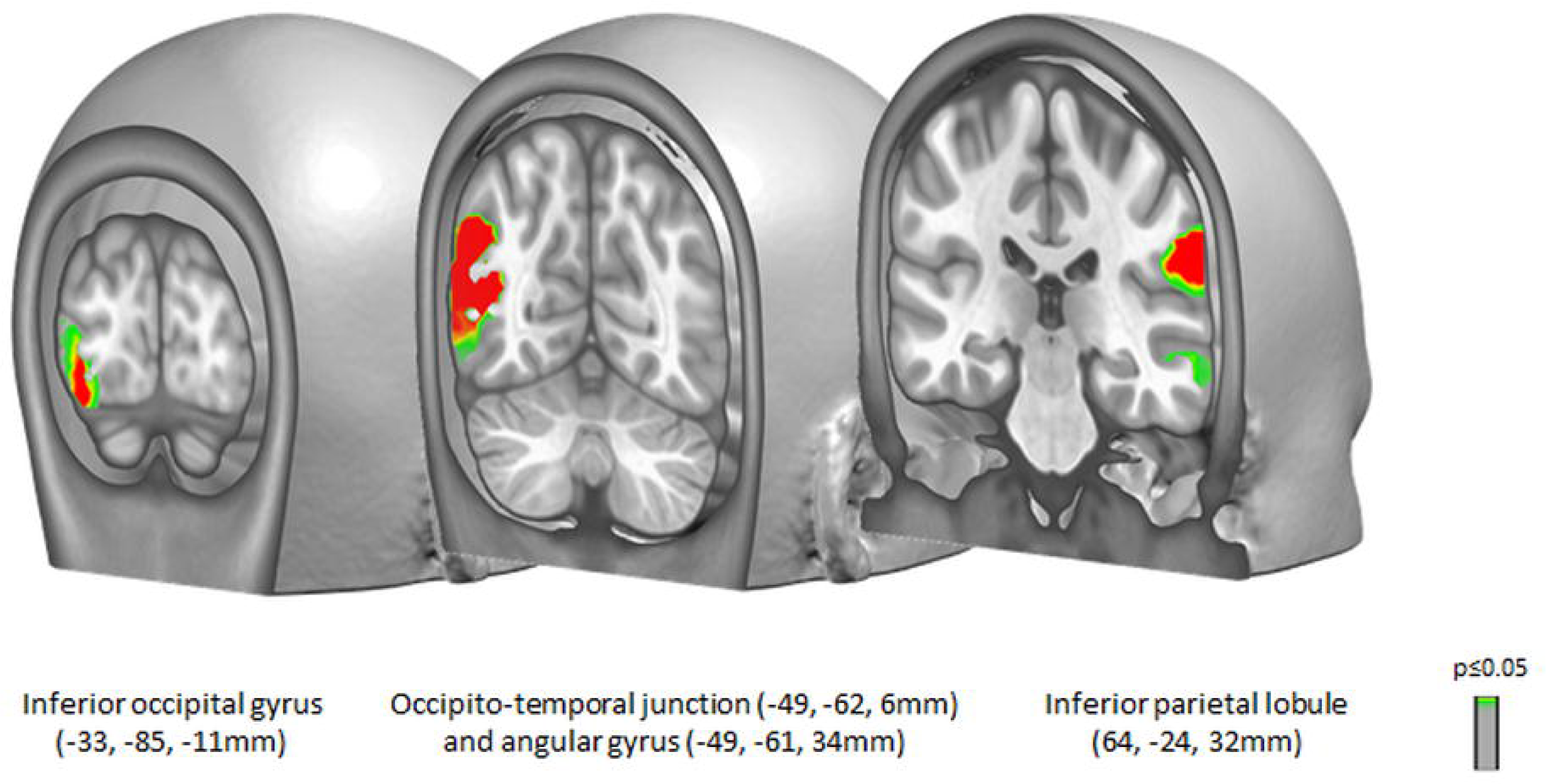
Source modelling analyses for responses to soundscapes of vertical and horizontal letters. Data from the 370-520ms post-sound onset time were then submitted to distributed linear source modelling. Stronger responses (p<0.05 for >10 contiguous solution points) to vertical letters were observed in the left occipital pole, the left occipito-temporal junction and angular gyrus; the Talairach^32^ coordinates of the local maxima are indicated. Stronger responses to horizontal letters were also observed in the inferior parietal lobe.

## Discussion

Our study provides the first characterization of the brain dynamics of SSD-mediated face and letter recognition skills in the blind. While canonical auditory cortex responses to SSD-soundscapes emerged at early and stereotypical latencies, they did not differentiate according to visual category information. In contrast, the first response that differentiated based on visually-conveyed information for both categories emerged in the visual cortices of the blind and were dissociable in both their timing and localization as a function of object category. ERPs to soundscapes of faces differed from those to scrambled faces within the first 460ms post-soundscape onset and followed from differential activation of brain networks that were localized to the FFA, lateral occipital cortices, and anterior temporal cortices. ERPs to soundscapes of vertically oriented letters differed from those to horizontally oriented letters within he first 370ms post-soundscape onset and followed from differential activation of brain networks principally within the left occipital pole as well as the left occipito-temporal junction extending to the angular gyrus. The spatio-temporal brain dynamics, afforded by electrical neuroimaging analyses of our EEG data, has allowed us to provide the key missing evidence that an image-to-sound SSD indeed is processed first and primarily in visual cortices. More generally, our findings uniquely reinforce the extant evidence for the profoundly task-contingent nature of the brain.

Image-to-sound sensory substitution has been shown to enable the recognition of a wide range of object categories from simple shapes through letters to faces and facial emotional expressions and body shape^4,6,12,15,33,34^. The hemodynamic imaging research, which hitherto dominated this research area, have provided an aggregate purview of the brain networks involved in SSD perceptions. Prior fMRI works have shown that responses in the VWFA are observed after, but not before, learning to identify the SSD shapes as letters (cf. Figure 4 in^6^). Other studies have documented sensitivity to SSD-conveyed body shapes within extrastriate body areas in the absence of evidence for sensitivity to such in auditory cortices of the superior temporal cortex^33^. However, despite subjective reports of the qualia of the substitution-mediated visual skills^35–37^, hemodynamic imaging research cannot provide direct evidence for whether those visual skills first differentially engage brain areas traditionally considered as visual, or if the latter are activated only as part of a widespread network of brain regions and subsequently to differentiate occurring elsewhere and/or earlier. Dynamic causal modelling has been applied to EEG data from blindfolded sighted participants who learned to use an SSD^38^. These results support a prominent role wherein visual mental imagery rather than cross-modal interactions. However, the conducted analyses did not characterize either the temporal dynamics (i.e., only oscillatory responses in a priori defined regions of interest were analyzed) or the capacity to discriminate specific categories of soundscapes (i.e., the data were collapsed across all correctly perceived stimuli). Thus, despite the presence of EEG findings in the area prior to the present study, the precise spatio-temporal brain dynamics of soundscape discrimination remained unknown.

Our study leveraged a combination of fine temporal resolution of EEG with the high spatial resolution of distributed EEG source modelling (collectively referred to as electrical neuroimaging^18–20,39^) to provide key missing evidence regarding the brain dynamics of SSD-conveyed face and letter perception. We capitalized upon the well-understood nature of brain sensitivity to faces and letter strings. The brain processes images of faces through a network of regions centered around the fusiform gyrus^21,22,40^ and lateral occipital face areas^41^. Our source localization results situate the earliest discrimination – i.e. starting at 460ms after soundscape onset - between soundscapes of faces and soundscape of scrambled faces precisely within this network of fusiform and lateral occipital cortices (**Figure 5**). Similarly, images of letters and words are processed starting at 370ms after soundscape onset in brain networks centered on the left fusiform gyrus (oftentimes referred to as the visual word-form area or VWFA^23^) as well as the angular gyrus^24^. While we cannot assert unequivocally whether our source localization included the VWFA, it is perhaps noteworthy that our local maximum in the occipito-temporal junction was situated <9mm from the centroid of the VWFA reported in the meta-analysis of Cohen et al.^42^ (see also^6^). Aside from letter processing in and of itself, another possibility is that the differential responses to the soundscapes of vertically and horizontally oriented letters reflects preserved sensitivity to orientation information in the congenitally blind.

It is worthwhile to situate the timing of the substitution-mediated face processing observed here alongside the more general findings of auditory object discrimination, as observed in sighted adults. Prior works have characterized a spatio-temporal hierarchy whereby animacy is discriminated within the initial 70-100ms of brain responses^43^, followed by discrimination of conspecific vocalizations at ∼170–220ms and at ∼290–360ms^44,45^ in turn followed by discrimination of action-related vs. action-free sounds of objects, at ∼300-360ms^46^ (reviewed in^47^). Additional work using single-trial EEG classification has shown there to be at least two phases of brain activity involved in perceptual decision-making regarding sound animacy: the first phase, spanning ∼110–180ms and is independent of perceptual capabilities, and the second phase, at ∼270–350ms, that manifests only during accurate categorizations^48^. The latency of our effects, while slightly later, indicates that also in our study, more complex perceptual decision-making processes had to be engaged, to distinguish faces from scrambled faces. In support with this idea, a study of Philiastides et al.^49^, involving perceptual decision-making with visual stimuli, has shown activations only at ∼450ms, when the task conditions were challenging. It is likewise important to consider that the soundscape presentation itself necessitates a temporal unfurling of the stimulus (i.e. the soundscape is rasterized from left to right). As such, the latency of the first effects indicative of sensitivity to soundscapes of faces at 460ms post-sound onset is not unreasonably “late” and is well within the idea that rapidly activated, sensory representations underlie substitution-mediated object recognition skills. Moreover, this latency aligns with the results of an ERP study of the discrimination of soundscapes of geometric forms in the sighted, wherein the recognition effects were observed 420–480 post-soundscape onset for the participants who learnt the vOICe SSD^25^.

Our study has several potential limitations. We obtained the data from a small, predominantly female sample, which may potentially curtail the strength of our conclusions. This sample size limitation is offset by the distinct spatio-temporal patterns of effects observed for faces and letters, respectively, and their localization to well-characterized brain regions implicated in visual processing of faces and letters. In fact, fMRI data were separately acquired in subsequent acquisitions from many of these participants^12^. The analysis here of multiple object categories – namely faces, scrambled faces, and subsets of letters – provides some measure of internal control conditions insofar as the effects differed in their timing and localization. More generally, our study highlights the added value of deeper data in situations of special or clinical populations^50^. The fact that participants listened passively to the soundscapes might also be considered a potential shortcoming. However, the psychophysical data obtained during a separate session would indicate that participants were able to discriminate soundscapes of different stimulus categories at near-ceiling levels. Thus, the SSD-mediated skills can be regarded as successfully acquired in our study. Additionally, the use of a passive paradigm has removed the risk of contamination of the categorical brain responses of interest by (pre)motor-related activations, which would be especially likely given the relatively “late” latency of the present effects. Furthermore, the category of soundscapes was randomized across trials and multiple exemplars from the same category were presented. These paradigm features make it unlikely that the present effects follow from any potential differences in allocation of attention. Further, the timing and sources of our effects are not readily explained in terms of attention modulations, which have been reported to occur within the initial ∼400ms post-sound onset^51^. Another shortcoming of our study was that ERP data were acquired only after, but not before, training with the vOICe SSD. Ideally, the data would have been obtained prior to and during training to show that the discrimination we report was not pre-existing and followed from training. The practical considerations of such a design notwithstanding, recordings prior to and during training might provide insights on how this ability is procured (see ^25^ for an example in sighted participants). However, this point would not detract from the principal observation here, that visual cortices canonically involved in visual face and letter discrimination are the first areas to exhibit discrimination when stimuli are conveyed as soundscapes even after adequate training. That is, one might even speculate that over-training could shift SSD-induced perceptions into auditory cortices, rather than engaging nominally visual areas. This proposition is in fact evident in the findings of Graulty et al.^25^, where the effects at 150 – 210ms post-soundscape onset correlated with performance accuracy.

More generally, our findings contribute to the accumulating evidence that the brain is fundamentally a task-contingent machine wherein neural architecture is constructed to fulfill a set of functions, rather than being defined by its nominal input sensory modality^15,52^. Our confirmation of this hypothesis has far-reaching implications. It follows that even brains that have been devoid of canonical sensory experience necessary for the regular functional brain organization to develop can achieve functional organization comparable to that observed in sighted individuals. However, this may be the case only if the missing sensory information becomes substituted with similar/complementary information through another sense. This notion would suggest that many of the findings indicating mass plasticity and cortical repurposing reported in the brains of the congenitally blind^53^ do not preclude the recruitment of these same brain regions to perform complex object recognition based on SSD training (see ^16^ for discussion of these concurrent theories). In conclusion, the current findings build on the extant evidence from the multiple hemodynamic and the scarce EEG studies, while simultaneously extending them critically, to offer new evidence for the spatio-temporal brain dynamics of SSD-elicited object recognition skills and the ability of the blind to functionally use their visual cortices for visual-like processing.

## Acknowledgments

Portions of this work have been funded by The Swiss National Science Foundation (grant 169206 to MMM), a fellowship from the Merkin Graduate Fellowship to LS, an ERC Consolidator grant “NovelExperiSENSE” to AA, and an Israeli Science Foundation grant to AA. We wish to acknowledge the contributions of Dr. Shlomo Bentin to this work.

## Author contributions

PJM: Writing of first draft, data analysis, edits and revisions

LR: Data collection, edits and revisions

LS: Writing of first draft, data curation, data analysis, edits and revision

CR: Data curation, data analysis

DA: Data analysis

ESA: Edits and revisions

EAV: Data collection, edits and revisions

AA: Conceptualization, supervision of data collection, study design, edits and revisions, and funding acquisition.

MMM: Conceptualization, writing of first draft, data curation, data analysis, edits and revisions, funding acquisition.

## Inclusion and diversity statement

As this was a convenience sample of congenitally blind individuals, gender balance and ethics or other types of diversity were challenging to ensure. Nonetheless, we worked to achieve the best balance possible. We worked to ensure that the study questionnaires were prepared in an inclusive way. One or more of the authors of this paper self-identifies as an underrepresented ethnic minority in their field of research or within their geographical location. One or more of the authors of this paper self-identifies as a member of the LGBTQIA+ community. While citing references scientifically relevant for this work, we also actively worked to promote gender balance in our reference list.

## STAR Methods

### Participants

Nine congenitally blind individuals (7 women) participated in the study (mean±sem age = 33.2±3.2 years). **Table 1** outlines the demographics of the participants including age, aetiology of blindness, and prior experience with the soundscape apparatus. Participants were recruited for this experiment from the group of participants that underwent long-term training with the vOICe SSD^54^. For all participants, consent was obtained prior to testing. The procedure was evaluated and approved by the ethics committee of the Hebrew University of Jerusalem. Data acquisition was completed between January and June 2012.

**Table 1.** Demographic information for each of the nine individuals.

| *Sex* | *Age* | *Condition* | *Light perception* | *Age of blindness onset (years)* |
| --- | --- | --- | --- | --- |
| F | 30 | ROP | None | 0 |
| M | 37 | ROP | None | 0 |
| F | 23 | Retinal detachment | None | 0 |
| F | 23 | Microphthalmia, Retinal detachment | None | 1 |
| F | 29 | Microphthalmia | None | 0 |
| F | 38 | Congenital rubella | None | 0 |
| M | 54 | ROP | None | 0 |
| F | 35 | ROP | None | 0 |
| F | 30 | LCA | Faint | 0 |

ROP = retinopathy of prematurity; LCA = Leber congenital amaurosis

### Procedure & stimuli

Participants were trained on a visual-to-auditory SSD termed the “vOICe”^17^. This device converts images into sounds, wherein position along the y-axis is translated into sound pitch (low to high), position along the x-axis is translated into time (left to right equals earlier to later in the sound sweep), and brightness is translated into volume. As the images here were black-and-white, there was no variation in volume. Example images used to generate soundscapes are shown in Figure 1. For more details on the training procedures and protocols, see ^54^. Before participating in the EEG portion of the study, each individual had completed over 70 hours of training with the vOICe over a period of several months. They then completed the EEG portion of the study with no overt task required. The EEG portion was comprised of 4 blocks of 60 trials each. In total, each participant was exposed to 240 soundscapes (i.e., 3 categories × 10 exemplars per category × 8 repetitions of each soundscape). Each soundscape had a duration of 1570ms. The inter-stimulus interval was comprised of 2 seconds of silence. During the EEG portion, soundscapes were heard passively. After the EEG, they then completed a 3-AFC discrimination task to ensure that each participant could reliably differentiate soundscapes of faces, scrambled faces, and Hebrew letters. This behavioural task entailed 90 trials (3 categories × 10 exemplars × 3 repetitions of each stimulus). Data from two participants on the behavioural task were corrupted and therefore could not be analysed. The performance of the remaining 7 individuals indicated near-ceiling accuracy (mean±s.e.m. faces = 95.7±1.6%; scrambled faces = 97.1±1.1%; letters = 97.6±1.0%). Reaction times on correct trials differed across conditions (faces = 1079±109ms; scrambled faces = 842±90ms; letters = 958±72ms) and were faster for soundscapes of scrambled faces than faces, though no other contrast was statistically significant. These reaction time data underscore the fact that soundscape stimuli, unlike their visual counterparts, do not appear in their entirely simultaneously, but rather unfurl over time.

To guide the analysis of ERPs to letter stimuli, we conducted a separate survey with 16 sighted adults none of whom were Hebrew speakers who rated each of the 10 visual letter stimuli on a −5 to 5 scale, with one extreme indicating strongly horizontal and the other extreme strongly vertical. The median ratings are shown in **Figure S1** and served as the basis for sub-grouping the letters into the 5 stimuli ranked as vertical and the 5 others ranked and horizontal/neutral. In this way, we retained the maximal numbers of EEG epochs.

### EEG recording and pre-processing

Continuous EEG was acquired at 256Hz through a 64-channel Biosemi ActiveTwo AD-box (www.biosemi.com), referenced to the common mode sense (CMS, active electrode) and grounded to the driven right leg (DRL; passive electrode), which functions as a feedback loop driving the average potential across the electrode montage to the amplifier zero. Data preprocessing and analyses were performed using the Cartool freeware (version 5.01^55^). Prior to epoching, the EEG was filtered (low-pass 40Hz; high-pass 0.1Hz; removed DC; 50Hz notch; using a second-order Butterworth filter with −12dB/octave roll-off that was computed linearly in both forward and backward directions to eliminate phase shifts). Peri-stimulus epochs, spanning 100ms pre-stimulus to 900ms post-stimulus onset, were averaged from each subject for each condition to compute event-related potentials (ERPs). Epochs were rejected based on automated artefact rejection criterion of ±100μV as well visual inspection for eye blinks and movement or other sources of transient noise.

The average (±SD) number of accepted EEG epochs for soundscapes of faces was 78.3 ± 2.06 for face (i.e., 98% of available trials) and for soundscapes of scrambled faces was 79.4 ± 1.33 (i.e., 99.8% of available trials) (mean±SD). These values did not significantly differ. Bad channels were identified before averaging and excluded from the artefact rejection. These data at artefact electrodes from each participant were interpolated using 3-D splines prior to group averaging^56^. The average number of interpolated channels was 0.25±0.44 (mean±SD). Additionally, we applied a pre-stimulus baseline correction and recalculated the data to the average reference.

For the analysis of responses to soundscapes of letters, the average number of accepted EEG epochs was 33±1.2 (i.e., 82% of the available trials) for the vertical condition and 32±2.7 (i.e., 80% of the available trials) for the non-vertical condition (hereafter, vertical and horizontal, respectively). These values did not differ significantly. As above, interpolation was performed using 3D-splines. On average, 1±1 channels were interpolated. Pre-stimulus baseline correction and average referencing were applied as above.

### ERP analyses design

Differences in the ERP responses to soundscapes of faces and scrambled faces as well as vertically and horizontally oriented letters were assessed using a multi-step analysis procedure, referred to as electrical neuroimaging, which involves global, reference-independent measures of the electric field at the scalp. These methods have been described in detail previously^18,19,57,58^.

The first reference-independent multivariate metric we analysed was the global field power (GFP), which is the spatial standard deviation across the electrode montage^59,60^. A stronger GFP value is indicative of greater and/or more synchronised brain activity, though the root cause (increased neural firing rate, increased numbers of active neurons, etc.) cannot be unequivocally asserted based on this measure alone. However, a modulation of GFP in the absence of reliable evidence for topographic modulations can most parsimoniously be interpreted as a modulation in the strength of responses originating from statistically indistinguishable sources or set of sources. The GFP was statistically analysed using a non-parametric paired randomization (1000 shuffles for each datapoint). An effect was required to last >10 continuous datapoints to be considered reliable and to account for temporal autocorrelation^26^.

The second reference-independent multivariate metric we analysed was the global map dissimilarity (GMD). This is the root mean square of the difference between two strength-normalized vectors (i.e. maps). In this way, GMD is exclusively sensitive to changes in topography. The GMD is at the core of data-driven topographic clustering^18^. This clustering was performed on the group-averaged data from both conditions. It is done to identify stable electric field topographies, referred to as “template maps”, and their pattern both in time and across conditions. The clustering is insensitive to pure amplitude modulations across conditions as the data are first normalised by their instantaneous GFP. The optimal number of template maps that explained the whole group-averaged data set was determined using a meta-criterion^61^. The clustering makes no assumption regarding the orthogonality of the derived template maps^62,63^. The pattern of maps identified in the group-average ERPs is used as a hypothesis generation tool that was then tested on single-subject data. was then submitted to a fitting procedure wherein each time point of each individual participant’s ERP is labelled according to the template map with which it best correlated spatially^18^. This yields a measure of relative map presence over a fixed time window (expressed as a percentage) that was then submitted to non-parametric analyses (specific test depending on the number of template maps back-fitted). This fitting procedure revealed whether responses to one type of soundscape were more often described by one ERP topography versus another, and therefore whether different intracranial generator configurations better accounted for responses to particular soundscapes. That is, biophysical laws indicate that a change in the topography of the electric field at the scalp surface forcibly originate from changes in the configuration of the underlying intracranial sources^19^.

Finally, we modelled the underlying intracranial sources of the ERPs in response to the two different types of soundscapes using a distributed linear inverse solution (minimum norm) combined with the LAURA (local autoregressive average) regularisation approach^64,65^. The solution space was calculated on a realistic head model that included 3000 nodes, selected from a grid equally distributed within the grey matter of the Montreal Neurological Institute’s average brain (available from https://github.com/DenisBrunet/Cartool). The head model and lead field matrix were generated with the Spherical Model with Anatomical Constraints (SMAC^66^ as implemented in Cartool version 5.01^55^, using a 6-shell model (CSF, skull compact, skull spongy, skull compact and scalp layers) as well as with an upper skull thickness (5.75mm) and mean conductivity (0.017S/m) values based on the mean age of our sample. As an output, LAURA provides current density measures; their scalar values were evaluated at each solution point. Statistical analysis of source estimations was performed by first averaging the ERPs across time for each participant and condition over the time window exhibiting a stable topography, described above. A non-parametric paired randomisation test was conducted across all solution points (1000 shuffles). The statistical significance criterion at an individual solution point was set at p<.0.05. Only clusters with ≥10 contiguous significant solution points were considered reliable in an effort to correct for multiple comparisons and was based on randomization thresholds (see also^29,31,48,67–69^ for similar implementations).

## Resource availability

### Lead contact

Further information and requests for resources should be directed to and will be fulfilled by the lead contacts, Micah Murray and Amir Amedi.

## Material, data and code availability

The stimuli, raw data and processed EEG data will be made available upon reasonable requests. The data were analysed using a freely available, GUI-based software for EEG analyses Cartool (https://github.com/DenisBrunet/Cartool).

