## Supplementary Figures for "The brain dynamics of congenitally blind people seeing faces and letters via sound"

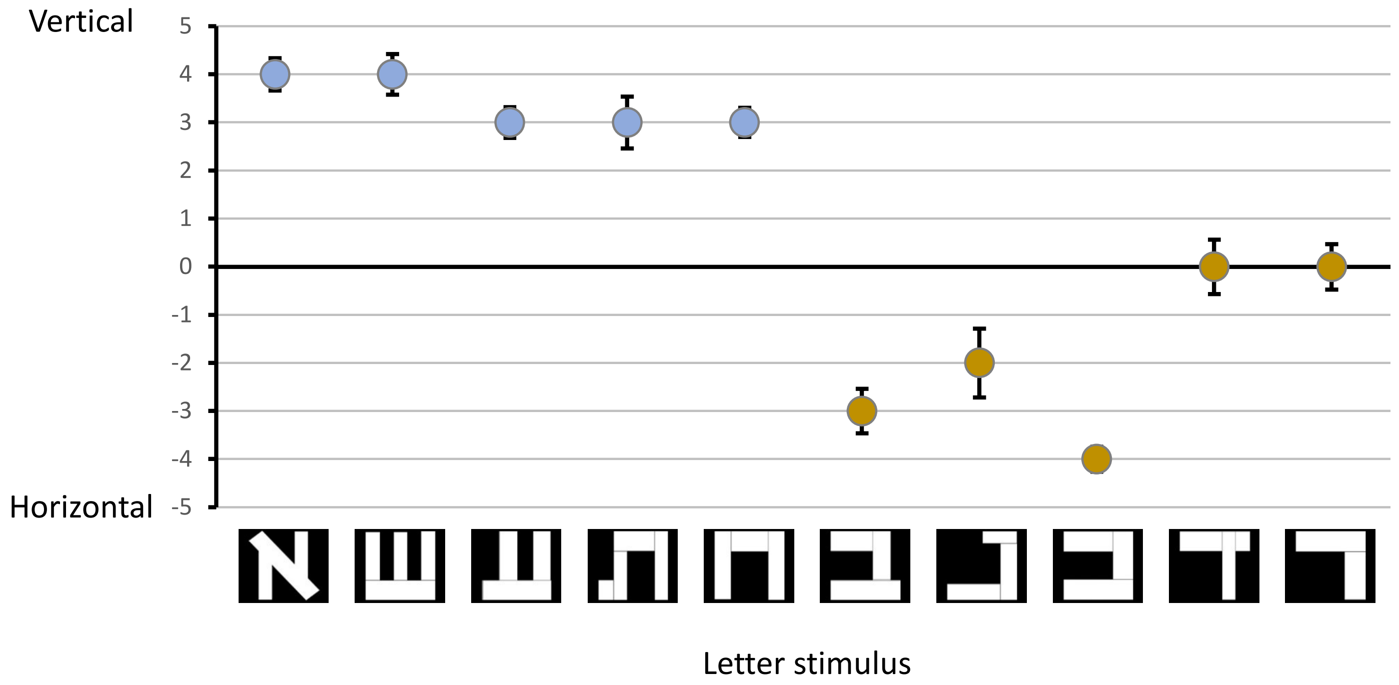


**Supplementary Figure S1**

### Single-subject back-fitting (720-900ms)

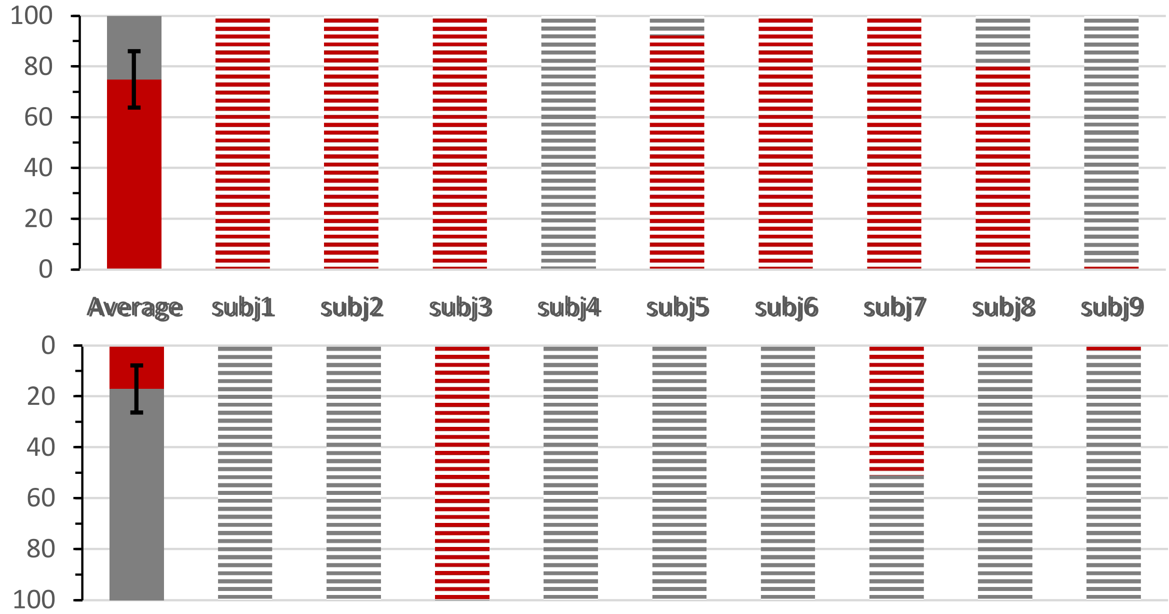


**Supplementary Figure S2**

### Single-subject back-fitting (524-900ms)

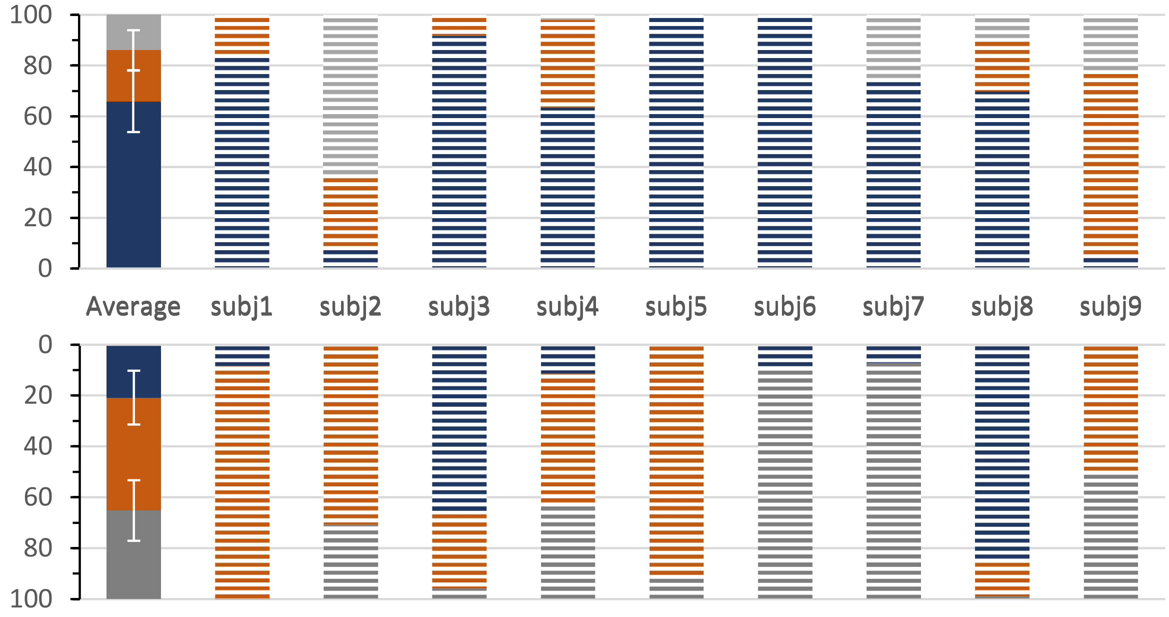


**Supplementary Figure S3**
